# Capture of fusion-intermediate conformations of SARS-CoV-2 spike requires receptor binding and cleavage at either the S1/S2 or S2’ site

**DOI:** 10.1101/2024.12.05.627124

**Authors:** Sabrina Lusvarghi, Russell Vassell, Brittany Williams, Haseebullah Baha, Sabari Nath Neerukonda, Carol D. Weiss

## Abstract

Although the structures of pre- and post-fusion conformations of SARS-CoV-2 spikes have been solved by cryo-electron microscopy, the transient spike conformations that mediate virus fusion with host cell membranes remain poorly understood. In this study, we used a peptide fusion inhibitor corresponding to the heptad repeat 2 (HR2) in the S2 transmembrane subunit of the spike to investigate fusion-intermediate conformations that involve exposure of the highly conserved heptad repeat 1 (HR1). The HR2 peptide disrupts the assembly of the HR1 and HR2 regions of the spike, which form six-helix bundle during the transition to the post-fusion conformation. We show that binding of the spike S1 subunit to ACE2 is sufficient to trigger conformational changes that allow the peptide to capture a fusion-intermediate conformation of S2 and inhibit membrane fusion. When TMPRSS2 is also present, an S2’ fusion intermediate is captured though the proportion of the S2’ intermediate relative to the S2 intermediate is lower in Omicron variants than pre-Omicron variants. In spikes lacking the natural S1/S2 furin cleavage site, ACE2 binding alone is not sufficient for trapping fusion intermediates; however, co-expression of ACE2 and TMPRSS2 allows trapping of an S2’ intermediate. These results indicate that, in addition to ACE2 engagement, at least one spike cleavage is needed for unwinding S2 into an HR2-sensitive fusion-intermediate conformation. Our findings elucidate fusion-intermediate conformations of SARS-CoV-2 spike variants that expose conserved sites on spike that could be targeted by inhibitors or antibodies.

**Author summary:** The SARS-CoV-2 spike protein undergoes two proteolytic cleavages and major conformational changes that facilitate fusion between viral and host membranes during virus infection. Spike is cleaved to S1 and S2 subunits during biogenesis, and S2 is subsequently cleaved to S2’ as the virus enters host cells. While structures of pre-fusion and post-fusion spike conformations have been extensively studied, transient fusion-intermediate conformations during the fusion process are less well understood. Here, we use a peptide fusion inhibitor corresponding to a heptad repeat domain in the S2 subunit to investigate fusion-inducing conformational changes. During spike-mediated cell-cell fusion, we show that the peptide binds to spike only after spike engages ACE2 and is cleaved at the S1/S2, S2’, or both sites. Thus, S2 needs at least one cleavage to refold to a peptide-sensitive fusion intermediate. SARS-CoV-2 variants differed in the proportion of S2 and S2’ fusion intermediates captured after receptor binding, indicating that the virus has evolved not only to alter its entry pathway but also to modulate S2 unfolding. This work informs the development of antiviral strategies targeting conserved sites in fusion-intermediate conformations of spike and contributes more broadly to the understanding of the entry mechanisms of viral fusion proteins.

## Introduction

The trimeric spike of the severe acute respiratory syndrome coronavirus 2 (SARS-CoV-2) mediates binding to host cell receptors and fusion between the viral and host membranes. Each spike monomer consists of two subunits (S1 and S2) that are generated by cleavage at the furin cleavage site (FCS) during biogenesis [1]. The receptor binding domain (RBD) in the S1 subunit spontaneously samples up and down conformations until it is captured by the angiotensin-converting enzyme 2 (ACE2) receptor on the host cell membrane. This initiates a series of conformational changes and exposes a second cleavage site (S2’) in the S2 transmembrane subunit to proteases on target cells [2]. The transmembrane serine protease 2 (TMPRSS2) is the primary protease involved in S2’ cleavage, but other proteases can also cleave S2 [3–5]. In models of spike-mediated fusion, the S2’ subunit then refolds, forming extended pre-hairpin, fusion-intermediate structures. This change repositions the N-terminal fusion peptide in S2’ for insertion into the target cell membrane. While S2’ is anchored in both viral and target cell membranes, the C-terminal heptad repeat region (HR2) in the extended intermediate folds back on the N-terminal heptad repeat (HR1), forming a thermostable coiled-coil, six-helix bundle (6HB) post-fusion structure [2]. This hairpin-like refolding brings the viral and host membranes together, allowing pore formation and membrane fusion for delivery of the viral genome into the host cell [6]. Only a limited number of high-resolution structures of unmodified spikes are available [7, 8]. The spike’s ability to undergo extensive conformational changes render it intrinsically metastable, thus modifications that increase the stability of the pre-fusion conformation have been developed [9]. Pre-fusion stabilizing mutations, such as K986P and V987P, that prevent the extension of the HR1-central helix (CH) junction needed to reach the post-fusion conformation, have been invaluable in both vaccine development [2] and in high resolution structure studies of the pre-fusion and ACE2-bound conformations [8, 10–16]. However, these mutations have limited the study of the conformational diversity intrinsic to the spike, as well as the knowledge of the fusion-intermediate conformations. The stabilized spikes allow binding to ACE2 but hinder the transition towards the intermediate conformations. Thus, when the stabilized spikes are used in vaccine development, the antibodies raised are limited to the pre-fusion conformation. Many highly conserved regions of the spike become exposed in the intermediate conformations [17], though potent neutralizing antibodies against those regions are rare [18].

Synthetic peptides corresponding to the heptad repeat (HR) regions in viral fusion proteins have been shown to inhibit membrane fusion and virus entry of various enveloped viruses, including HIV and several coronaviruses [19–23]. HR peptide fusion inhibitors bind to the viral fusion proteins during conformational changes and interfere with formation of the 6HB needed for fusion in a dominant-negative manner. HR2 derived peptides were shown to be effective at inhibiting both SARS-CoV-2 infection and cell-cell fusion [24] and have emerged as potential candidate antivirals in animal models [25]. Lipid conjugation, dimerization, and hydrocarbon stapling of SARS-CoV-2 HR2 peptides enhance antiviral potency and in vivo half-life [25–29]. Analysis of the structure-activity relationship for SARS-CoV-2 HR2-derived peptides has elucidated the significance of N- and C-terminal amino acid residues [27, 28, 30]. Such peptides can also be valuable tools for investigating conformational changes. For instance, a dimeric cholesterol-HR2-derived peptide was employed in cryo-electron tomography (Cryo-ET) to trap and visualize an extended and partially folded intermediate [31].

In this study, we use an HR2-derived peptide to capture fusion-intermediate conformations of spike to dissect steps in the fusion process. We investigate peptide inhibition and binding to spike under various experimental conditions and compare differences among variants. Our work demonstrates how receptor activation and protease cleavages at S1/S2 and S2’ sites regulate S2 unwinding, as spike refolding facilitates membrane fusion.

## Results

### The inhibitory potency of the 36-HR2 peptide depends on the virus entry pathway

In selecting an HR2 peptide for our investigations, we first compared potency of two HR2 derived peptides, a 36-residue HR2 peptide (36-HR2) and an overlapping 42-residue HR2 peptide (42-HR2) with six more N-terminal residues (Figs 1A and 1B) that have been reported to have different potencies under different experimental conditions [27, 30]. We found that both 36-HR2 and 42-HR2 peptides comparably inhibited D614G (SARS-CoV-2 Wuhan variant with D614G mutation) lentiviral pseudovirus infections in 293T cells stably expressing both ACE2 and TMPRSS2 (ACE2/TMPRSS2 target cells), with half-maximal inhibitory concentration (IC_50_) in the low µM range (1.06 and 1.65 µM, respectively, S1A Fig). We chose to use the shorter version of the peptide (36-HR2) for our studies.

**Fig 1.**
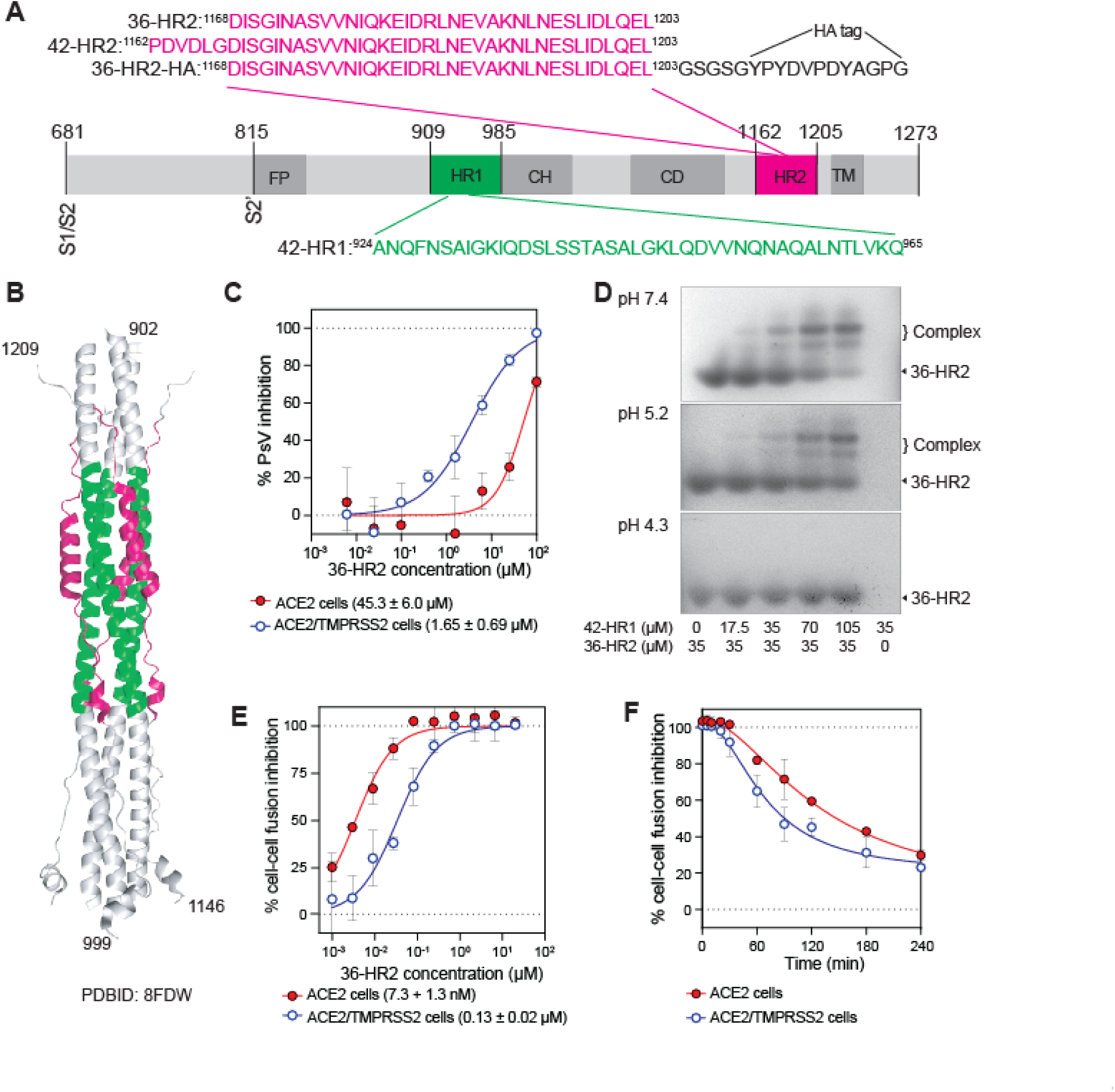
The inhibitory potency of the 36-HR2 peptide depends on the virus entry pathway. **(A)** Schematic representation of SARS-CoV-2 S2 protein and the HR1 (green) and HR2 (magenta) derived peptides. Functional domains of the S2 protein: FP, fusion peptide; HR1, heptad repeat 1; CH, central helix; CD, connector domain; HR2, heptad repeat 2; TM, transmembrane domain. The S1/S2 and S2’ cleavage sites are marked. **(B)** Cartoon representation of the 6-helix bundle region of the SARS-CoV-2 spike in the post-fusion conformation (PDB 8FDW). The 42-HR1 and 36-HR2 are highlighted in green and magenta respectively. **(C)** Dose-response curves of 36-HR2 peptide inhibition of D614G pseudovirus infection of 293T cells expressing ACE2 (red filled circles) or ACE2/TMPRSS2 (blue open circles). **(D)** Interaction of 42-HR1 and 36-HR2 peptides at different pHs. Bands corresponding to 36-HR2 and the 42-HR1+ 36-HR2 complexes are indicated. **(E)** 36-HR2 peptide inhibition of spike-mediated cell-cell fusion between 293T effector cells expressing D614G spike and 293T targets cells expressing ACE2 (filled red circles) or ACE2/TMPRSS2 (open blue circles). **(F)** Time-dependence of 36-HR2 inhibition of cell-cell fusion between D614G spike effector cells and ACE2 (red filled circles) or ACE2/TMPRSS2 target cells (blue open circles). In **(C)** and **(E)** each datapoint corresponds to the average of three independent experiments, and the mean IC_50_ and the standard deviation of three independent experiments is shown below each graph. In **(F)** one representative out of three experiments with similar results is shown.

Viral infection can occur by fusion with the plasma membrane or with endocytic vesicles [32]. It was previously shown that HR2-derived peptides inhibited SARS-CoV viral infection through the cell surface pathway more than through the endocytic pathway [21]. Thus, we compared the potency of the peptide against SARS-CoV-2 pseudoviruses in 293T cells that expressed only ACE2 (ACE2 target cells) or both ACE2 and TMPRSS2 (ACE2/TMPRSS2 target cells). We found that 36-HR2 was 27-fold more potent in ACE2/TMPRSS2 target cells than ACE2 target cells (Fig 1C). Previously we showed that our pseudoviruses use the endocytic entry pathway in these ACE2 target cells [33]. Several factors may account for the decreased peptide potency for viruses in the endocytic pathway, including lower pH and reduced peptide concentrations in the endosomes. To assess if acidification in the endosomes could contribute to the weaker potency of the peptide in ACE2 target cells, we investigated the effect of pH on the affinity of the 36-HR2 for its target HR1 domain. The HR1 and HR2 peptides form a complex that mimics the 6HB present in the post-fusion conformation of S2. We mixed the 36-HR2 with a 42 amino acid peptide corresponding to HR1 (42-HR1, Fig 1A, green) at different pH conditions and assessed complex formation by native gel shift experiments (Fig 1D). The 42-HR1 peptide isoelectric point (pI) is 8.54 and thus migrates in the opposite direction in native gel electrophoresis from the 36-HR2 and the 6HB complex (pI of 4.2 and 4.5, respectively). We observed two bands, likely corresponding to a five-helix bundle and the 6HB, as the proportion of 42-HR1 increased (Fig 1D). Complex formation decreased as the pH decreased, suggesting that protonation of the negatively charged residues in HR1 or 36-HR2 may hinder the complex formation and contribute to the decrease in peptide potency against virus entry in the endocytic pathway.

We next investigated peptide inhibition of spike-mediated cell-cell fusion. The peptide inhibited cell-cell fusion of spike-expressing 293T cells (effector cells) with ACE2 target cells in the low nanomolar range (Fig 1E, red circles, 7.3 ± 1.3 nM IC_50_) and with ACE2/TMPRSS2 target cells in sub-micromolar range (Fig 1E, open blue circles, 0.13 ± 0.02 µM IC_50_). Notably, the peptide more potently inhibited spike-mediated cell-cell fusion with ACE2 target cells than ACE2/TMPRSS2 target cells, in contrast to the pattern that was seen for virus-cell fusion (Figs 1C and 1E). This finding suggests that availability of TMPRSS2 or other cell surface proteases affects the rate of fusion and half-life of the peptide-sensitive conformations of spike.

To assess if the lower potency of the peptide in the presence of TMPRSS2 correlated with faster fusion and transition to the post-fusion conformation, we added 36-HR2 at different timepoints after mixing effector cells and target cells. We found that cell-cell fusion was sensitive to peptide inhibition for longer times in ACE2 target cells than in ACE2/TMPRSS2 target cells (Fig 1F), suggesting that TMPRSS2 accelerates the spike-mediated fusion, in agreement with a previous report [34]. Complementing other studies that highlight differences in viral fusion and cell-cell fusion [4, 35–37], our data shows that the availability or exposure of the intermediate conformations might differ in these systems.

We note that the peptide potency depended on the levels of spike expression in the effector cells. With high levels of spike expression (0.25-4 µg spike expression plasmid in transfections), the peptide potency decreased (26.5-274 nM in ACE2 target cells and 0.47-6.13 µM in ACE2/ TMPRSS2 cells, S1B Fig). Interestingly, at intermediate 36-HR2 concentrations (10-100 nM) the peptide enhanced the cell-cell fusion (S1B Fig). This enhancement, which was more pronounced in ACE2/TMPRSS2 target cells, may be due to sub-stoichiometric inhibition by peptide that may lead to stabilization of fusion-intermediate conformations in a proportion of spikes involved in a multi-spike fusion pore. An analogous inhibitor enhancement of SARS-CoV-2 infectivity has also been seen for soluble ACE2 (sACE2) [38].

### Spike binding to ACE2 allows the 36-HR2 peptide to capture an S2 fusion intermediate, while TMPRSS2 cleavage allows capture of an S2’ second fusion intermediate

To investigate the spike conformations captured by the peptide during membrane fusion, we used an HA epitope-tagged 36-HR2 peptide (36-HR2-HA) to bind to spike in flow cytometry and co-immunoprecipitation experiments. The HA tag did not alter the peptide potency (S1A Fig). In flow cytometry experiments with mixtures of effector cells expressing D614G spike and ACE2 or ACE2/TMPRSS2 target cells (Fig 2A), the peptide efficiently bound effector cells only in the presence of ACE2 or ACE2/TMPRSS2 target cells (Fig 2B, red and blue traces respectively). Control target cells with no ACE2 nor TMPRSS2 expression showed minimal peptide binding (Fig 2B, gray trace), suggesting that the HR1 region is mostly not available for peptide binding during spontaneous sampling or breathing of unbound spikes.

**Fig 2.**
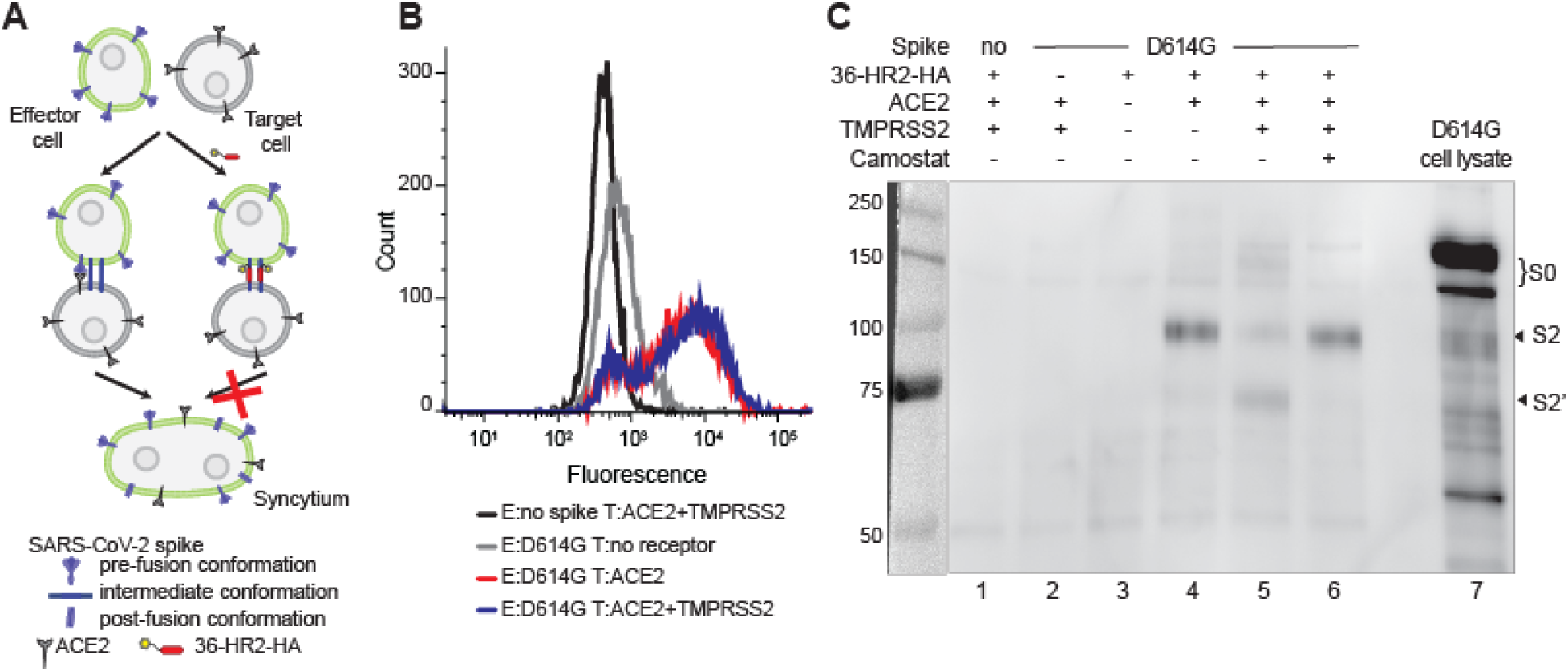
ACE2 binding is sufficient to allow peptide capture of an S2 fusion intermediate, while TMPRSS2 allows binding to an S2’ fusion intermediate. **(A)** Experimental setup to assess trapping of fusion-intermediate conformations of spike. In the absence of peptide, spike and ACE2 interact, favoring the conformational changes in spike that facilitate membrane fusion and syncytia formation. In the presence of peptide, the spike cannot transition to the post-fusion conformation and syncytia is not formed. Formation of the trapped intermediate conformations can be analyzed by flow cytometry on the cell surface or by co-immunoprecipitation on cell lysates. **(B)** Flow cytometry histograms showing the interaction of 36-HR2-HA with effector cells (E), expressing D614G and target cells (T), expressing ACE2 (red trace) or ACE2/TMPRSS2 (blue trace). Control effector cells lacking spike (black trace) and target cells lacking receptors (gray trace) are shown. After incubation of HA-tagged peptide with cells, cells were washed and then incubated first with anti-HA, followed by Alexa 680 anti-mouse antibodies. One representative out of three experiments with similar results is shown. **(C)** Capture of the D614G spike fusion intermediates by co-immunoprecipitation with 36-HR2 peptides. Effector cells expressing D614G spike were incubated with 36-HR2-HA in the absence of receptor or in the presence of ACE2 or ACE2/TMPRSS2 target cells in the absence or presence of camostat. Cells were washed and lysed after incubation with the peptide. Peptide-captured spike was immunoprecipitated with anti-HA conjugated beads. Immunoprecipitated samples were analyzed by western blotting using and anti-S2 antibody. The S2, S2’, and S0 bands are indicated.

To further elucidate the spike intermediates captured by the peptide, we performed co-immunoprecipitation experiments with the 36-HR2-HA peptide, similar to the capture of fusion-intermediate conformations of the HIV envelope glycoprotein [39–41]. The 36-HR2-HA was incubated with spike-expressing effector cells and ACE2 or ACE2/TMPRSS2 target cells. The 36- HR2-HA preferentially co-immunoprecipitated S2 in the presence of ACE2 target cells (Fig 2C, lane 4). A small amount of S2’ was also co-immunoprecipitated, likely resulting from low level S2’ cleavage by membrane metalloproteases. The proportion of S2’ that was co-immunoprecipitated increased in the presence of ACE2/TMPRSS2 target cells (Fig 2C, lane 5). Addition of camostat, a TMPRSS2 inhibitor, decreased the proportion of S2’ to similar levels with ACE2 target cells (Fig 2C, lane 6). Control experiments without peptide, effector cells without spike, or target cells without ACE2 or TMPRSS2 (Fig 2C lanes 1, 2 and 3 respectively) showed no S2 or S2’ bands. Efforts to perform analogous co-immunoprecipitation experiments with pseudovirus were not successful due to the low amounts of spike in the pseudoviruses.

### At least one spike cleavage is needed for unwinding of S2

Having established that the peptide binds to two forms of receptor-activated spike, we next investigated whether the S1/S2 cleavage was needed for trapping of these intermediate conformations. We engineered two furin cleavage site (FCS) mutants: D614G ΔFCS, which deletes residues 681-684 (PRRA), and D614G FCS(SKPSK), which replaces residues 682-684 (RRA) by a Factor XA cleavage site (Fig 3A). Factor XA was previously shown to cleave the S1/S2 in SARS-CoV [42]. Pseudovirus infection of both mutants was less dependent on the presence of TMPRSS2 (S2A Fig), consistent with previous reports that FCS deletion mutant viruses have less of a preference for the TMPRSS2 pathway than wild type [36, 43]. 36-HR2 inhibited infection of both pseudovirus mutants with comparable IC_50_ values as D614G (Fig 3B, IC_50_ values 5.60 ± 2.43 and 1.94 ± 0.68 µM for D614G ΔFCS and D614G FCS(SKPSK), respectively).

**Fig 3.**
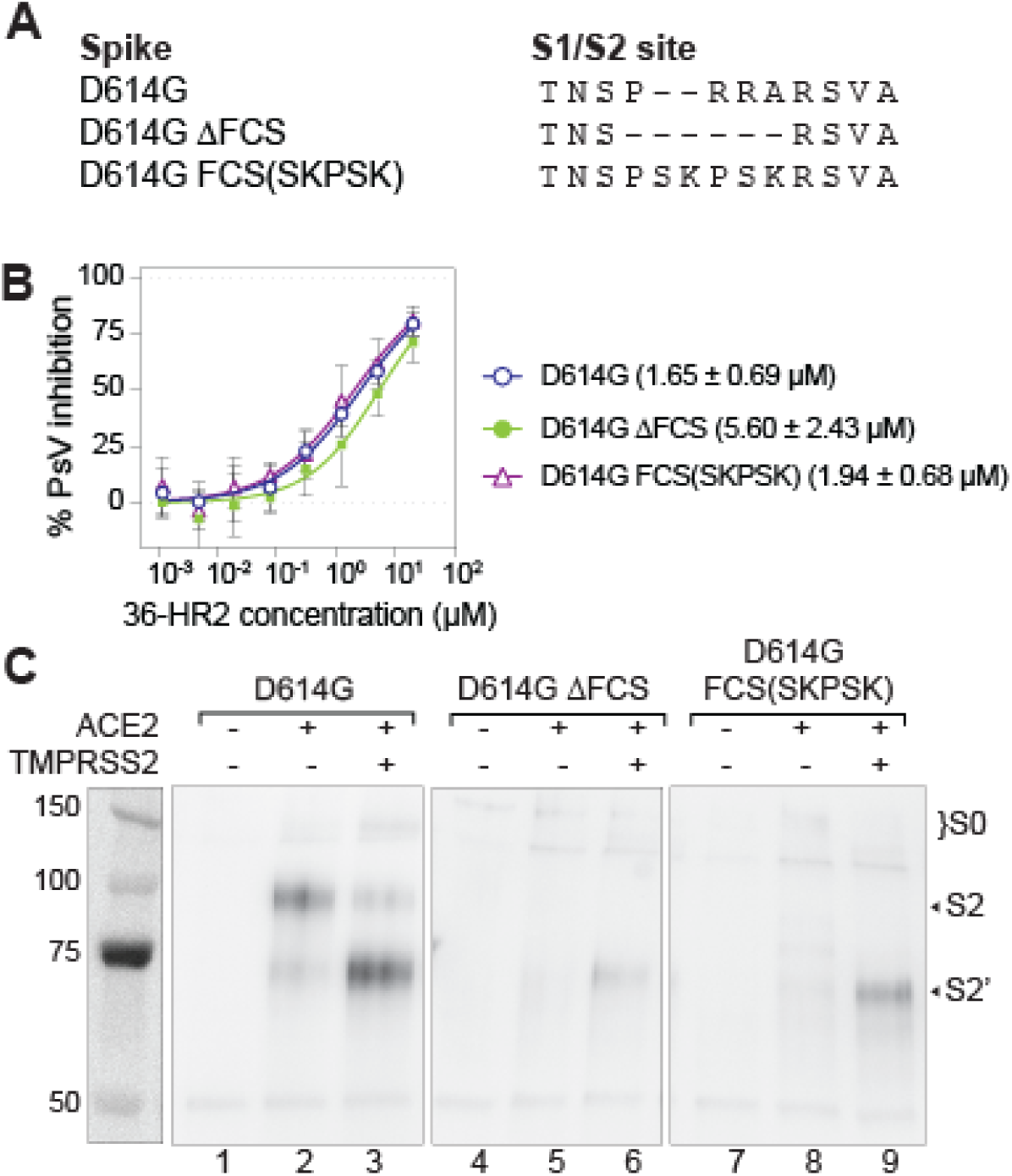
At least one spike cleavage is needed for unwinding of S2 to expose the HR1 in the fusion-intermediate conformation of spike. **(A)** Amino acid sequence alignment of the S1/S2 junction region with the furin cleavage site (FCS) of spikes used in this study. **(B)** 36-HR2 peptide inhibition of infection of ACE2/TMPRSS2 target cells by pseudoviruses with or without FCS spike mutations. Each datapoint corresponds to the average of three independent experiments. Mean IC_50_ and the standard deviation of three independent experiments is shown. **(C)** Co-immunoprecipitation of the D614G, D614G ΔFCS and D614G(SKPSK) spikes with 36-HR2-HA in the absence or the presence of ACE2 or ACE2/TMPRSS2 target cells. The S2, S2’, and S0 bands are indicated. One representative out of two experiments with similar results is shown.

In cell-cell fusion assays we found that both FCS spike mutants did not mediate significant cell-cell fusion with ACE2 target cells (S2B Fig), in agreement with prior reports showing that spike-mediated cell fusion is severely impaired in S1/S2 mutants in the absence of TMPRSS2 [4, 5, 36, 44]. Accordingly, the peptide failed to capture full length (S0) or S2 fusion intermediates for these mutants in co-immunoprecipitation experiments (Fig 3C, lanes 5 and 8). Addition of TMPRSS2 in the target cells rescued cell-cell fusion to some extent (S2B Fig, 6.3- and 3.0-fold increase for D614G ΔFCS and D614G FCS(SKPSK), respectively). In this case, the peptide co-immunoprecipitated a band corresponding to S2’ in both D614G ΔFCS and D614G FCS(SKPSK) spikes (Fig 3C, lanes 6 and 9).

### Some RBD binding antibodies trigger the fusion-intermediate conformation

Next, we used the peptide to explore how RBD-binding antibodies might affect spike conformational changes. We investigated CB6 (a precursor of etesevimab) and bebtelovimab (Beb) monoclonal antibodies because they both overlap the ACE2 binding site but recognize spike conformations differently. Due to steric hindrance, ACE2 and CB6 can only bind the RBD in the “up” conformation [45, 46] (Fig 4A, blue and green respectively). Beb binds RBD in a region that overlaps the ACE2 binding site, which is accessible in both “up” and “down” conformations [47] (Fig 4A, red). Both antibodies inhibited spike mediated cell-cell fusion with ACE2/TMPRSS2 target cells with IC_50_s in the low µg/mL range (Fig 4B). To assess if these antibodies could trigger the spike and allow exposure of the HR1 region, effector cells expressing D614G spike were pre-treated with antibody for 30 minutes before the addition of 36-HR2 and target cells expressing no receptor, ACE2, or ACE2/TMPRSS2. In the presence of CB6, peptide-trapped S2 was observed regardless of the presence of ACE2 or TMPRSS2 in target cells (Fig 4C, lanes 6-8). These data show that CB6, which traps the “up” conformation of the RBD, triggers the formation of a 36- HR2 sensitive, S2 intermediate. Further, TMPRSS2 cleavage was diminished when CB6 was used as a trigger, highlighting the role of ACE2 in keeping the spike near the membrane for TMPRSS2 cleavage. By contrast, preincubation of the Beb antibody with the effector cells completely prevented the peptide trapping of either the S2, or S2’ intermediate with ACE2 or ACE2/TMPRSS2 target cells (Fig 4C, lanes 11 and 12, respectively). Although structural studies of Beb binding to the RBD or stabilized spike show that Beb binds to the RBD in closed and open conformations [47], our findings show that in contrast to CB6, Beb binding to functional spike prevents conformational changes involving exposure of the HR1 region.

**Fig 4.**
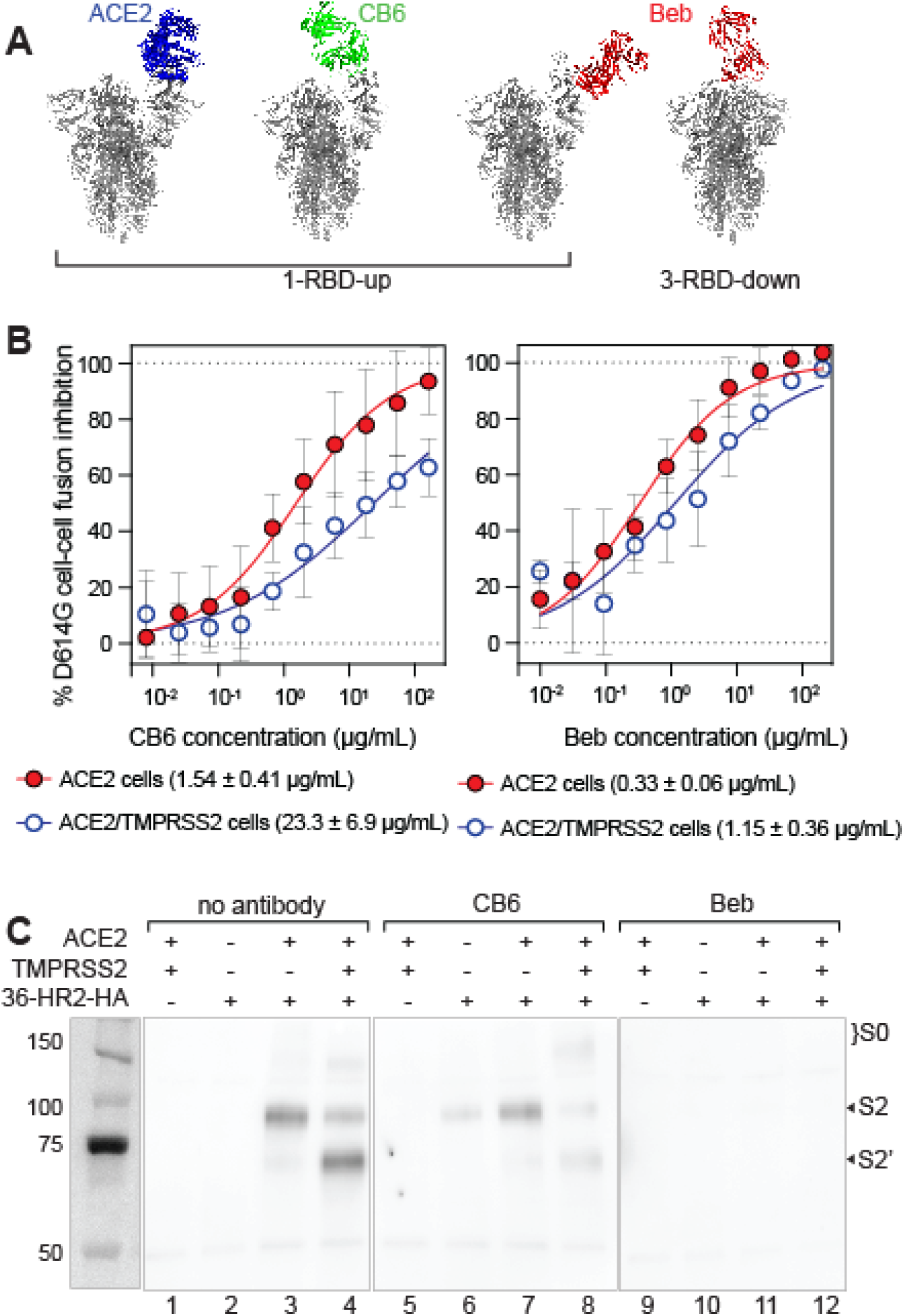
Binding of the CB6 antibody to the RBD triggers spike conformational changes to fusion intermediates. **(A)** Superimposition of the structures of the SARS-CoV-2 spike trimer with 1-RBD-up conformation (PDB 7KDJ, gray) and structures of the RBD complexed with ACE2 (PDB 7A94, blue) and CB6 (PDB 7C01, green). Superposition of the 1-RBD-up and 3-RBD-down conformations (PDBs 7KDJ and KDG, respectively, gray) and the structure of the RBD complexed with Beb (PDB 7MMO, red). **(B)** Inhibition of CB6 and Beb against D614G spike-mediated cell fusion with 293T cells expressing ACE2 (red filled circles) or ACE2/TMPRSS2 (blue open circles). One representative out of three experiments with similar results is shown. Mean IC_50_ and the standard deviation of three independent experiments is shown. **(C)** Antibody bound spikes were co-immunoprecipitated with 36-HR2-HA in the absence or the presence of ACE2 or ACE2/TMPRSS2. Effector cells were pre-incubated with CB6 or Beb antibodies before the co-immunoprecipitation. The S2, S2’ and S0 bands are indicated. One representative out of two experiments with similar results is shown.

### Variants differ in the proportion of S2 and S2’ fusion intermediates that are trapped by the HR2 peptide

We next examined whether the spikes of the Wuhan-Hu-1 (WA1), D614G, Delta, BA.1, and XBB.1.5 variants differed in their transition to fusion-intermediate conformations of spike. The HR2 domain is conserved among these variants, but the HR1 domain has substitutions in Delta (D950N), BA1 (Q954H, N969K and L981F), and XBB.1.5 (Q954H and N969K) (Fig 5A and 5B). We note that transfecting equal amounts of spike plasmids resulted in different levels of expression of the spike variants (S3A Fig), though all variants similarly induced syncytia in the presence or absence of TMPRSS2 (S3B Fig). We found that the 36-HR2-HA peptide potently inhibited cell-cell fusion of these variants and the potency against all variants was lower with ACE2/TMPRSS2 than ACE2 target cells, as was the case for the D614G variant (S3C Fig). Difficulties in controlling comparable spike expression levels among these variants precluded a direct comparison of peptide potency against these variants in the cell-cell fusion assay.

**Fig 5.**
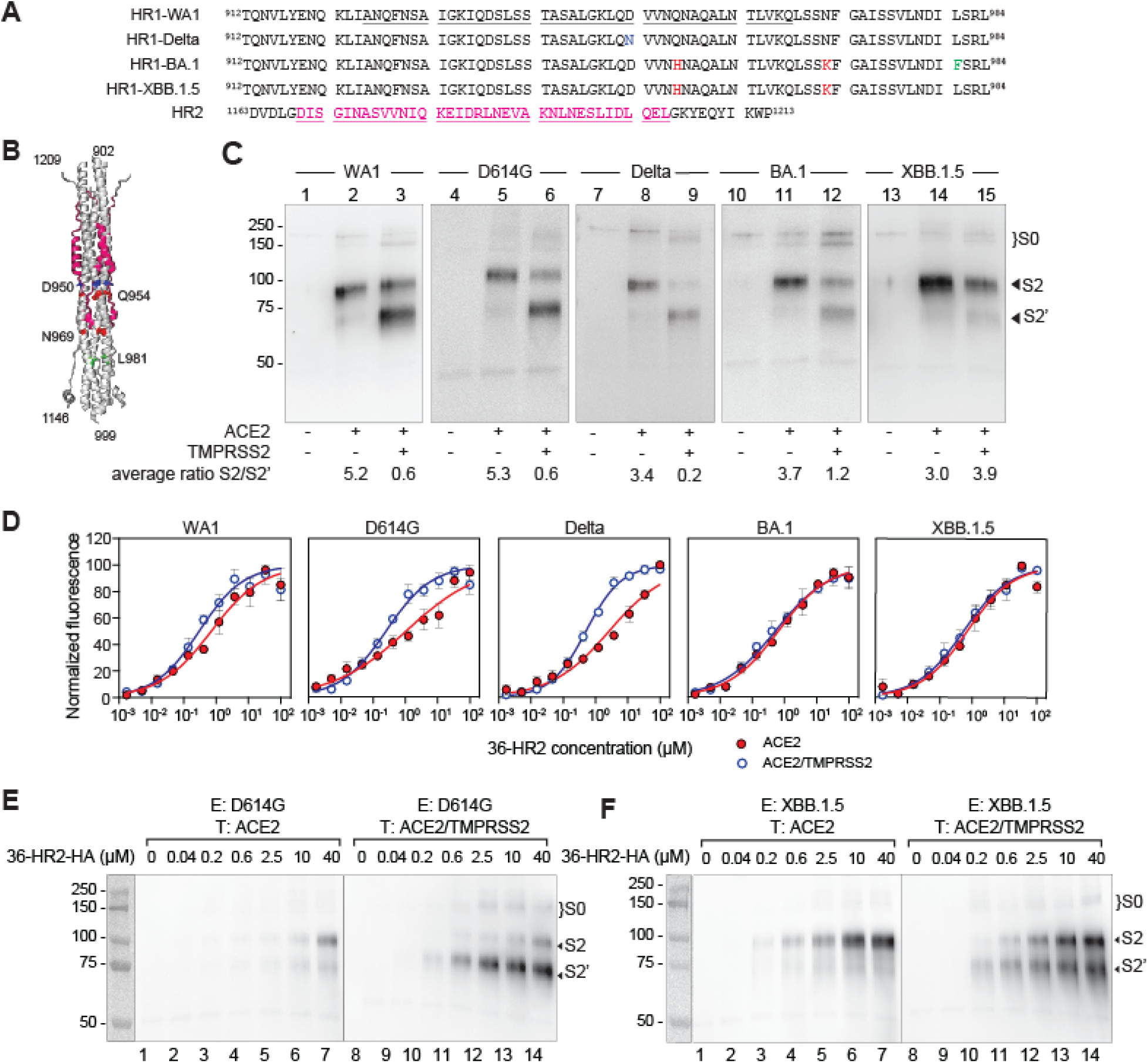
Trapping of the S2 intermediate, rather than S2’ is favored in Omicron variants in the presence of TMPRSS2. **(A)** Amino acid sequence alignment of the HR1 region in different mutants used in this study. Mutations are color coded. The sequence of the HR2 region is also presented. 42-HR1 and 36-HR2 peptides are underlined. **(B)** Cartoon representation of the 6-helix bundle region of the SARS-CoV-2 post-fusion conformation (PDB 8FDW). The 36-HR2 is highlighted in magenta. Residues mutated in Delta and Omicron variant are highlighted. **(C)** Spikes corresponding to the different variants were co-immunoprecipitated with 36-HR2-HA in the absence of any receptor, or in the presence of ACE2 or ACE2/TMPRSS2. Samples from the co-immunoprecipitation were analyzed by western blot using and anti-S2 antibody. The S2, S2’ and S0 bands are indicated in each panel. S2/S2’ densitometry ratio is presented below each lane. **(D)** 36-HR2 binding to different variants against 293T cells expressing ACE2 (red circles) or ACE2/TMPRSS2 (open blue circles) as measured by flow cytometry. One representative out of three experiments with similar results is shown. Characterization of the D614G **(E)** and XBB.1.5 spike **(F)** intermediates by co-immunoprecipitation as a function of peptide concentration. Effector cells, E, expressing D614G or XBB.1.5 spike were incubated with variable amounts of 36-HR2- HA with target cells, T, expressing ACE2 or ACE2/TMPRSS2. After incubation cells were washed and lysed. Anti-HA labeled beads were used to co-immunoprecipate the peptide and the spikes. Samples were then analyzed by western blot using and anti-S2 antibody. The S2, S2’ and S0 bands are indicated in each panel.

In peptide co-immunoprecipitation experiments, addition of ACE2 target cells was sufficient to trigger spike conformational changes that allowed peptide trapping of S2 fusion intermediates in all variants (Fig 5C lanes 2, 5, 8, 11 and 14). With ACE2/TMPRSS2 target cells, different proportions of the S2 and S2’ fusion intermediates were captured among the variants. The proportion of peptide-captured, S2’ fusion-intermediate compared to the S2 fusion intermediate was highest for the Delta spike, medium for the D614G and WA1 spikes, and lowest for the BA.1 and XBB.1.5 spikes (Fig 5C, lanes 3, 6, 9, 12 and 15). These findings are consistent with the relative efficiency of spike cleavage by TMPRSS2 among these variants and their preferences for the endocytic entry pathways [48–50].

To further investigate whether the 36-HR2-HA peptide preferentially binds to the S2 or S2’ fusion intermediate, we performed peptide dose-response experiments in flow cytometry and co-immunoprecipitation studies. In flow cytometry experiments, the binding of 36-HR2 to D614G and Delta spikes was stronger with ACE2/TMPRSS2 target cells than ACE2 target cells (Fig 5D, 4.1 and 5.3-fold, respectively), while peptide binding to WA1, BA.1 and XBB.1.5 spikes showed no differences between ACE2 and ACE2/TMPRSS2 target cells (Fig 5D, 2.3, 1.2 and 1.3-fold respectively). In peptide dose-response, co-immunoprecipitation experiments, we investigated D614G and the most antigenically distant XBB.1.5 variant. We found that both D614G and XBB.1.5, allowed S2 trapping by 36-HR2-HA in the presence of ACE2 target cells in a dose-dependent manner (Figs 5E and 5F, respectively, lanes 1-7). In ACE2/TMPRSS2 cells, the S2’ was captured by peptide at lower peptide concentrations than S2 for the D614G spike, suggesting stronger affinity for the S2’ intermediate (Fig 5E, lanes 8-14). This result indicates that the HR1 region may become more accessible upon the cleavage of the S2’ site. For the XBB.1.5 spike, lower concentrations of peptide preferentially captured more S2’ than S2 with ACE2/TMPRSS2 target cells, but higher concentrations favored S2 trapping (Fig 5F, lanes 8-14).

## Discussion

Here, we used an HR2 peptide to elucidate fusion-intermediate conformations of spike that involve unwinding of the S2 subunit during spike-mediated membrane fusion. We demonstrate that spike binding to ACE2 or the CB6 antibody that stabilizes the up conformation of the RBD is sufficient to trigger unwinding of S2 and expose the HR1 region. While buried regions such as the fusion peptide, have been shown to become accessible even without ACE2 engagement [51]—indicating that spike breathing is not limited to the movement of the RBD—details about the intermediate conformations in unmodified spikes remain unresolved despite computational studies combined with mass spectrometry studies [52–54]. Our findings offer new details about S2 transitions that complement structural snapshots of spike in pre-fusion, receptor-bound, and post-fusion conformations.

Our co-immunoprecipitation data revealed that the spike transitions through at least two distinct fusion intermediates, S2 and S2’, in the process of mediating membrane fusion. The first intermediate, S2, can be trapped by the 36-HR2 after ACE2 engagement, while TMPRSS2 cleavage facilitates trapping of a second intermediate, S2’ (Fig 6, step 3). Both intermediates may involve formation of a 3-helix, HR1 coiled coil that could be stabilized by interaction with the 36- HR2 [55], leading to formation of a stable 6HB (Fig 6, step 4). Although both fusion intermediates can be trapped in different variants, the relative proportion of S2 and S2’ intermediates captured was lower in Omicron spikes, in agreement with inefficient processing of S2 to S2’ at the cell surface [49].

**Fig 6.**
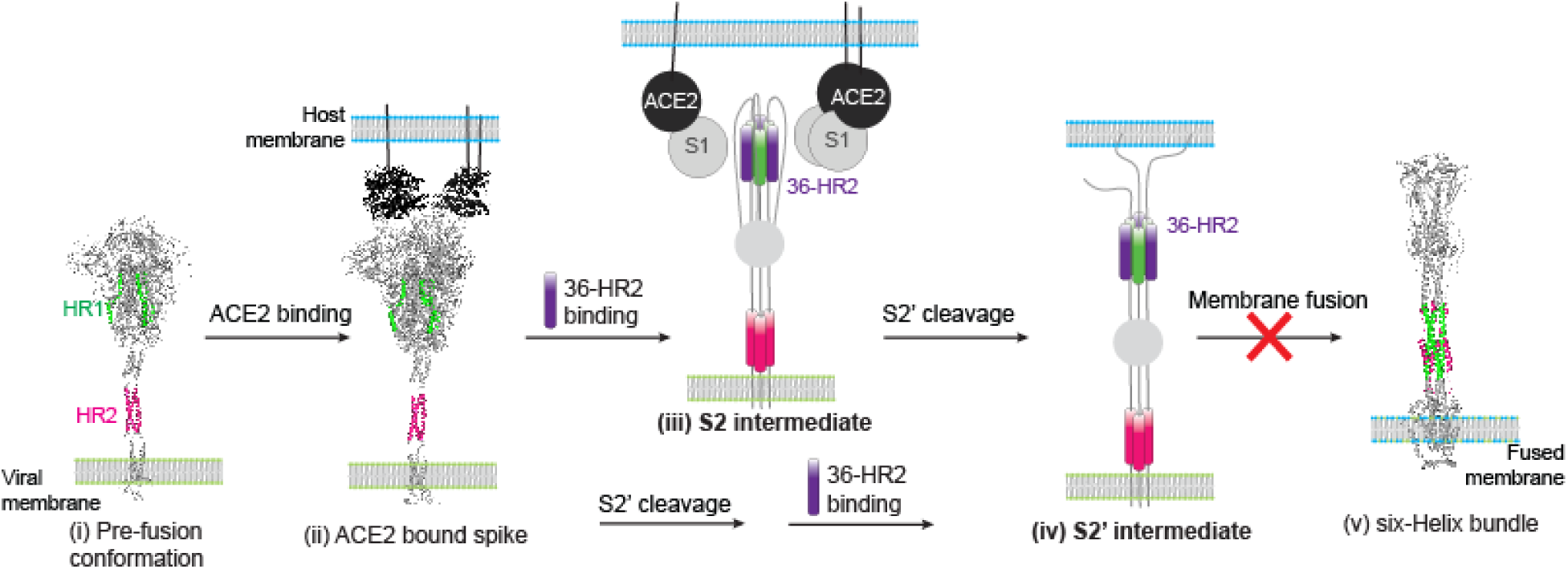
Model of spike-mediated membrane fusion showing capture of 36-HR2/S2 fusionintermediates. Schematic of viral entry of spike-mediated membrane fusion. Full length model of the spike glycoprotein from ref [56] was used to highlight the HR1 (green) or HR2 (magenta) regions in the pre-fusion conformation. In the pre-fusion conformation (i), the RBD continuously samples the up and down conformations until it binds to ACE2 to form an ACE2-spike complex (ii). ACE2 binding facilitates an open conformation that exposes the HR1 to 36-HR2 (purple) binding, trapping the S2 intermediate (iii). Formation of this intermediate may concur with S1 subunit displacement. The S2’ cleavage site is exposed to proteolysis by TMPRSS2 or other metalloproteases. A second S2’ intermediate is trapped by 36-HR2 (iv). Trapping of the S2’ intermediate may occur before or after S2’ cleavage. The binding of 36-HR2 to HR1 during transition to the fusion intermediate interferes with formation of the spike six-helix bundle (v) preventing membrane fusion.

Notably, ACE2 binding alone was sufficient for trapping the S2 intermediate, but only if spike had been cleaved at the S1/S2 junction. In the absence of a FCS, cleavage at S2’ was required for transitioning to a peptide-sensitive fusion intermediate. Thus at least one spike cleavage is required for S2 unwinding to reach HR1-accessible fusion intermediates. Our dose-response experiments suggest that TMPRSS2 processing of spike to generate S2’ improves peptide binding (Fig 5E and 5F), though we cannot rule out the possibility that the peptide-captured S2 intermediates are subsequently cleaved by TMPRSS2.

The rate of spike-induced membrane fusion can also modulate the ability of the 36-HR2 peptide to capture fusion intermediates. The lower potency of the peptide in inhibiting cell-cell fusion with ACE2/TMPRSS2 target cells than ACE2 target cells correlated with a shorter time to transition to the peptide-insensitive, post-fusion conformation (Fig 1E and 1F). These findings may be explained by less efficient cleavage of spike to generate S2’ by alternate serine-proteases when TMPRSS2 is absent. Subsequent transition of spike to the post-fusion conformation likely results in longer exposure of the peptide-sensitive conformations.

In contrast to cell-cell fusion, the potency of the peptide was greater with ACE2/TMPRSS2 than ACE2 target cells. Lower concentrations of peptide in the endosomes, reduced peptide binding to the HR1 under low pH conditions, faster triggering of the spike at low pH, or a combination of these and other factors may all contribute to the lower potency of peptide inhibition of pseudovirus infection via the endocytic pathway. Paradoxically, we also found that the peptide can enhance viral infection at certain concentrations (S1B Fig). Examples of concentration-dependent, dual effect compounds have been reported, including physiological concentrations of recombinant ACE2 [38, 57]. This consideration is important when designing and evaluating to use peptides fusion inhibitors and vaccines that target fusion intermediates.

The potency of peptide inhibition may also depend on the dynamics of fusion pore formation. Spike-mediated membrane fusion likely involves not only S2 refolding in a single spike trimer but also coordinated refolding of multiple spike trimers to achieve the post-fusion conformation needed to create a fusion pore. Slower pore formation could extend the half-life of HR2 peptide-sensitive spikes. A combination of factors, including spike density, receptor availability, and exposure of the S2’ cleavage site, may influence the number of spikes in intermediate conformations waiting to synchronously transition to the post-fusion conformation. The number of spike trimers needed to form a fusion pore for virus entry or cell-cell fusion could also differ and affect peptide potency. In the case of HIV, it has been shown that the number of spikes needed for fusion is variant dependent [58], with some variants requiring a single spike for fusion [59].

Overall, our results highlight spike transitions in the S2 subunit requiring spike proteolytic processing and ACE2 engagement. These findings refine our understanding of the spike entry mechanism that offer insights into the development of inhibitors and vaccines as well as the evolution of spike in different variants. Limitations in our study include the use of spike expression plasmids and cell culture conditions that may not completely mimic authentic virus infection and cell-cell fusion *in vivo*.

## Materials and methods

### Plasmids and cell lines

VRC8400 or pcDNA3.1(+) plasmids were used for expression of codon optimized WA1 (Wuhan strain), D614G (WA1 + D614G), Delta, BA.1 or XBB.1.5 SARS-CoV-2 spikes. Spike-plasmids were purchased from GenScript, (Piscataway, NJ). HIV gag/pol (pCMVΔR8.2), luciferase reporter (pHR’CMV-Luc), human ACE2 (hACE2-TM) and TMPRSS2 (pCAGS- TMPRSS2) were obtained from the Vaccine Research Center (VRC), National Institute of Allergy and Infectious Diseases (NIAID), National Institutes of Health (NIH), Bethesda, MD. pSCTZ- alpha plasmid and Omega cells were kindly provided by Nathaniel Landau, (New York University). pCAGGS-TMPRSS2 plasmid was kindly provided by Dr. Mikhail Matrosovich (University of Marburg, Germany). HEK293-T-ACE2 (293T.ACE2s) cells stably expressing ACE2 were obtained through BEI Resources, NIAID, NIH; (NR-52511, contributed by Jesse Bloom, Fred Hutchinson Cancer Research Center, Seattle, WA) [60]. 293T-ACE2/TMPRSS2 target cells stably expressing ACE2/TMPRSS2 were established and maintained as previously described [33]. Cell lines were cultured in Dulbecco’s modified essential medium (DMEM) supplemented with penicillin/streptomycin, non-essential amino acids, L-glutamine, HEPES and 10% fetal bovine serum at 37 °C with 5% CO2. Omega cells were grown in the same conditions but in the presence of 0.2 mg/mL hygromycin.

### Peptides and other reagents

The 36-HR2, 42-HR2, 36-HR2-HA and 42-HR1 peptides were synthesized and purified by the CBER Facility for Biotechnology Resources at the FDA. HR2-peptide stocks were prepared in phosphate buffered saline (PBS) at 100-400 µM. 42-HR1 peptide stock was prepared in water at 100-400 µM. Only for the experiment comparing the potency of 36-HR2 against D614G pseudovirus in ACE2 and ACE2/TMPRSS2 target cells, which required higher concentrations, a peptide stock of 4.8 mM was prepared in DMSO.

The CB6 monoclonal antibody was generated by transfecting 3.60 x 10^8^ 293-Freestyle cells (ThermoFisher Scientific, Waltman, MA) in FreeStyle 293 media (ThermoFisher Scientific) with heavy and light chain expression plasmids (kindly provided by Peter Kwong, Vaccine Research Center, NIH) using 293fectin (ThermoFisher Scientific) transfection reagent. Cultures were incubated at 37°C with 8% CO_2_ at 125 rpm and harvested at 72 hours post transfection. Cells were removed by centrifugation and the supernatant was filtered with a 0.45 μm filter followed by a 0.20 μm filter. CB6 was purified from the supernatant using a Cytiva HiTrap Protein-G column, buffer exchanged to DPBS and concentrated using an Amicon-15 Ultra centrifugal filter with 30 kDa molecular weight cut off. Bebtelovimab was generously provided by Eli Lilly (Indianapolis, IN). Camostat mesylate was purchased from Millipore-Sigma (Burlington, MA).

### SARS-CoV-2 pseudovirus production and peptide inhibition assays

Pseudovirus production: HIV-based lentiviral pseudoviruses were produced by co-transfecting 4 µg of spike pcDNA3.1(+) or 0.5 µg of spike VRC8400 plasmids, 5 µg of the pCMVΔR8.2 (HIV gag/pol) plasmid, and 5 µg of the pHR’CMV-Luc (luciferase reporter) plasmid in 10-cm dishes containing 50-80% confluent 293T cells in 10% FBS/DMEM using Fugene 6 (Promega, Madison, WI). 48 h after transfections, pseudoviruses were harvested and supernatants were filtered through 0.45 µm low protein binding filters and aliquoted at −80°C until further use.

Peptide inhibition assays: Pseudoviruses with titers 0.8-1.5^6^ relative luminescence units (RLU)/mL were incubated with 4-fold serial dilutions of peptide for 1.5 h at 37°C. The peptide-virus mixtures (100 µL) were added to ACE2 or ACE2/TMPRSS2 target cells that were seeded in 96-well plates at a density of 0.3×10^6^ cells/mL (100 µL/well) 24 hours earlier. Pseudovirus infection was allowed to continue for 48/72h. Pseudovirus infectivity was quantified by luciferase activity (luciferase assay system, Promega, Madison, WI). Inhibition curves were normalized to pseudovirus only control and fitted using nonlinear regression curve [inhibitor] vs normalized response (GraphPad Prism, La Jolla, CA). The peptide concentration corresponding to 50% neutralization was defined as IC_50_. All experiments were performed in at least two independent experiments, each with an internal intra-assay replicate.

### SARS-CoV-2 Spike-mediated cell-cell fusion inhibition assay

Preparation of effector and target cells: Effector cells were prepared in 10-cm dishes containing 50-80% confluent 293T cells in 10% FBS/DMEM and were co-transfected with 0.05 µg (unless specified) of spike-pcDNA3.1(+) or spike-VRC8400 plasmids and 5 µg of pSCTZ- alpha plasmid using Fugene 6. Target cells were prepared in 10-cm dishes containing 50-80% confluent Omega cells in 10% FBS/DMEM and were co-transfected with 2 µg of ACE2 plasmid and 1 µg of empty pcDNA3.1(+) (ACE2 target cells) or 2 µg of ACE2 plasmid and 1 µg of TMPRSS2 plasmid (ACE2/TMPRSS2 target cells) using Fugene 6.

Cell-cell fusion inhibition: 24h after transfection, cells were harvested using cell dissociation solution (Sigma Aldrich) and resuspended at 0.6×10^6^ cells/mL. 50 µL aliquots of 4x serial dilution of peptide (or antibody) and 50 µL effector cells were combined followed by addition of 50 µL target cells. Cells were co-cultured for 18-22h at 37°C. Syncytia formation was quantified by beta-galactosidase activity (Galacto-star™, ThermoFisher Scientific). Effector cells expressing the alpha subunit but no spikes and target cells expressing no ACE2 or TMPRSS2 were used to assess the background activity. Neutralization curves were normalized to the beta-galactosidase activity in effector cells and target cells combined without addition of any peptide and fitted using nonlinear regression curve [inhibitor] vs normalized response (GraphPad Prism, La Jolla, CA). The peptide concentration dilution corresponding to 50% neutralization was defined as IC_50_. All experiments were done at least in duplicates, each an internal intra-assay replicate.

Time dependent addition of 36-HR2: 50 µL aliquots of effector and target cells were combined. 50 µL of 36-HR2 (13.3 µM final concentration) was added at different time points (0- 240 min). Cells were co-cultured for 18-22h at 37°C and syncytia formation was quantified by beta-galactosidase activity (Galacto-star™, ThermoFisher Scientific). Effector cells expressing spikes and target cells expressing empty pcDNA3.1(+) plasmid were used to assess the background activity. Neutralization curves were normalized to the beta-galactosidase activity in effector cells and target cells combined without addition of any peptide and plotted as a function of time of addition (GraphPad Prism, La Jolla, CA). All experiments were done at least in duplicates, each with an internal intra-assay replicate.

### Co-immunoprecipitation assay

Preparation of effector and target cells: 10-cm dishes containing 50-80% confluent 293T cells in 10% FBS/DMEM were co-transfected as follows, effector cells were transfected with 4 µg of spike pcDNA3.1(+) or 1 µg spike VRC8400 plasmids. Target cells were transfected with 2 µg of ACE2 plasmid (ACE2 only cells) or 2 µg ACE2 and 1 µg of TMPRSS2 plasmids (ACE2/TMPRSS2 target cells). Empty pcDNA3.1(+) was co-transfected in all plates to complete 6 µg total plasmid. After 24h cells were harvested using cell dissociation solution (Sigma Aldrich) and resuspended at 10×10^6^ cells/mL. For co-immunoprecipitation a total of 5×10^6^ cells/mL effector cells, 36-HR2-HA (40 µM final concentration, unless otherwise stated), camostat (0 or 500 µM final concentration) and 5×10^6^ cells/mL target cells were mixed in 1.5 ml microcentrifuge tubes. The mixture was twirled for 2h at 37°C. Cells were washed two times with media, then lysed overnight using 1% NP-40 buffer. The clarified lysate was allowed to bind anti-HA agarose beads (Sigma Aldrich) for 30 min at room temperature. Beads were washed 4 times with 1% NP-40 buffer then resuspended in Laemmli sample buffer (Bio-Rad) containing 10 µM DTT and subjected to denaturing electrophoresis in 10% Mini-PROTEAN TGX gels (Bio-Rad). Proteins were then transferred to a nitrocellulose membrane (Bio-Rad) and membranes were blocked overnight with 2% BSA (Sigma-Aldrich). Membranes were probed with anti-S2 (Genetex), washed, probed with anti-mouse-IgG-HRP (Seracare), washed, and bands detected with Lumiglo-Reserve (Seracare) on a G:Box (Syngene).

Experiments to assess the binding of peptide in the presence of antibody were performed as described above but bebteovimab or CB6 were added to the effector cells and incubated for 30 minutes before addition of peptide and target cells.

### Flow cytometry assay

10-cm dishes containing 50-80% confluent 293T cells in 10% FBS/DMEM were co-transfected with 4 µg of spike pcDNA3.1(+) or 1 µg spike VRC8400 plasmids and 0.5 µg of eGFP plasmid (Addgene plasmid # 160697) [61] (Effector cells) or 2 µg of ACE2 plasmid (ACE2 target cells) or 2 µg ACE2 and 1 µg of TMPRSS2 plasmids (ACE2/TMPRSS2 target cells). Empty pcDNA3.1(+) was co-transfected in all plates to complete equal amounts of total plasmid. After 24h cells were harvested using cell dissociation solution (Sigma Aldrich) and resuspended at 10×10^6^ cells/mL. A total of 0.5×10^6^ cells/mL effector cells, 36-HR2-HA (40 µM final concentration, unless otherwise stated), and 0.5×10^6^ cells/mL target cells were combined and incubated for 2h at 37°C. Cells were spun at 2000g and then washed 2x with flow cytometry buffer (FCB, PBS + 2% FBS + 0.5% Sodium azide). Anti-HA antibody (Bioxcell, 025 mg/mL, 100 µL) in FCB was allowed to bind for 30 min at room temperature. Cells were washed 2x with FCB, and TRITC Goat Anti-Mouse IgG (H+L) (Jackson ImmunoResearch, 1:150 dilution, 100 µL) in FCB was allowed to bind for 30 min at room temperature. Cells were washed twice then fixed using 2% formaldehyde in FCB and stored for 16-40h before flow cytometry analysis. Samples were analyzed using a BD LSR Fortessa Cell Analyzer (BD Biosciences), and FlowJo 10.10.0 software was used to process the flow cytometry data.

### Gel-shift assay

The 36-HR2 peptide (25 µM) was combined with the 42-HR1 peptide (0-75 µM) for 30 min at 37°C, with PBS (pH 7.4), 10 mM Sodium Acetate (pH 5.2) or 10 mM Citric Acid (pH 4.3). 50% glycerol was the added to the samples (5% final concentration). Samples were subjected to native gel electrophoresis in 10% Mini-PROTEAN TGX gels (Bio-Rad). Proteins were stained using InstantBlue Coomassie Protein Stain (Abcam).

## Author contributions

Conceptualization SL, RV, CW

Formal Analysis SL, CW

Funding Acquisition CW

Investigation SL, RV, BW, HB

Methodology SL, RV, SN

Project Administration SL, CW

Resources CW

Supervision SL, CW

Visualization SL, CW

Writing-Original Draft Preparation SL, CW

Writing-Review and Editing SL, RV, BW, HB, SN, CW

## Acknowledgments

We thank Hugo Azurmendi and Jason Gorman (U.S. Food and Drug Administration) for critical reading of the manuscript. We also thank the U.S. Department of Health and Human Services, Office of the Assistant Secretary for Preparedness as part of the U.S. Government COVID-19 response Therapeutics Research Team for providing reagents for this project.

## Supporting information

**S1 Fig. Characterization of the inhibitory activity of HR2 peptides in pseudovirus infection and cell-cell fusion.**

**(A)** Inhibition of pseudovirus infectivity by 36-HR2 (black circles), 42-HR2 (green circles) and 36-HR2-HA (pink circles) against D614G pseudovirus infection of 293T cells expressing ACE2/TMPRSS2. **(B)** Inhibition activity of D614G spike mediated cell fusion with 293T cells expressing ACE2 (filled circles) or ACE2/TMPRSS2 (open circles) by 36-HR2.

**S2 Fig. FCS spike mediated pseudovirus infectivity and cell-cell fusion.**

**(A)** Infectivity of pseudoviruses bearing FCS modified spikes of target cells expressing ACE2 (green) and ACE2 and TMPRSS2 (red). The data shows averaged relative luminescence units (R.L.U.), error lines represent the standard deviations of three independent experiments. Ratio between the infectivity for each pseudotypes in ACE2 and ACE2/TMPRSS2 are presented above each pair (p≤0.05, *; p≤0.01, **, Wilcoxon matched-pairs signed rank test). **(B)** Spike mediated cell fusion effector cells expressing D614G or the FCS modified spikes and target cells expressing ACE2 or ACE2/TMPRSS2. The data shows averaged relative luminescence units (R.L.U.), error bars represent the standard deviations of three independent experiments (p>0.05, ns; p≤0.05, *, One-Way ANOVA, Tukey’s multiple comparisons test with a single pooled variance).

**S3 Fig. 36-HR2 inhibition of spike-mediated cell-cell fusion of different SARS-CoV-2 variants.**

**(A)** Western blot showing expression levels of different spikes in the effector cells. **(B)** Spike mediated cell fusion effector cells expressing spike corresponding to different variants (WA1, D614G, Delta, BA.1 and XBB.1.5) and target cells expressing ACE2 or ACE2/TMPRSS2. The data shows averaged relative luminescence units (R.L.U.), error bars represent the standard deviations of three independent experiments (p>0.05, ns; p≤0.05, *; p≤0.01 **; p≤0.001, ***, One-Way ANOVA, Tukey’s multiple comparisons test with a single pooled variance). **(C)** 36-HR2 inhibition of cell-cell fusion of different variants with 293T cells expressing ACE2 (red circles) or ACE2/TMPRSS2 (blue triangles). Each datapoint corresponds to the mean and the standard deviation of at least three independent experiments.

**S4 File. Data file.**

## References

1. Chan YA, Zhan SH. The Emergence of the Spike Furin Cleavage Site in SARS-CoV-2. Mol Biol Evol. 2022;39(1). doi: 10.1093/molbev/msab327. PubMed PMID: 34788836; PubMed Central PMCID: PMCPMC8689951.

2. Jackson CB, Farzan M, Chen B, Choe H. Mechanisms of SARS-CoV-2 entry into cells. Nat Rev Mol Cell Biol. 2022;23(1):3–20. Epub 20211005. doi: 10.1038/s41580-021-00418-x. PubMed PMID: 34611326; PubMed Central PMCID: PMCPMC8491763.

3. Simmons G, Bertram S, Glowacka I, Steffen I, Chaipan C, Agudelo J, et al. Different host cell proteases activate the SARS-coronavirus spike-protein for cell-cell and virus-cell fusion. Virology. 2011;413(2):265–74. Epub 20110323. doi: 10.1016/j.virol.2011.02.020. PubMed PMID: 21435673; PubMed Central PMCID: PMCPMC3086175.

4. Hornich BF, Grosskopf AK, Schlagowski S, Tenbusch M, Kleine-Weber H, Neipel F, et al. SARS- CoV-2 and SARS-CoV Spike-Mediated Cell-Cell Fusion Differ in Their Requirements for Receptor Expression and Proteolytic Activation. J Virol. 2021;95(9):e00002–21. Epub 20210412. doi: 10.1128/JVI.00002-21. PubMed PMID: 33608407; PubMed Central PMCID: PMCPMC8104116.

5. Nguyen HT, Zhang S, Wang Q, Anang S, Wang J, Ding H, et al. Spike glycoprotein and host cell determinants of SARS-CoV-2 entry and cytopathic effects. J Virol. 2021;95(5):e02304–20. Epub 20201211. doi: 10.1128/JVI.02304-20. PubMed PMID: 33310888; PubMed Central PMCID: PMCPMC8092844.

6. Kreutzberger AJB, Sanyal A, Saminathan A, Bloyet LM, Stumpf S, Liu Z, et al. SARS-CoV-2 requires acidic pH to infect cells. Proc Natl Acad Sci U S A. 2022;119(38):e2209514119. Epub 20220901. doi: 10.1073/pnas.2209514119. PubMed PMID: 36048924; PubMed Central PMCID: PMCPMC9499588.

7. Cai Y, Zhang J, Xiao T, Peng H, Sterling SM, Walsh RM, Jr., et al. Distinct conformational states of SARS-CoV-2 spike protein. Science. 2020;369(6511):1586–92. Epub 20200721. doi: 10.1126/science.abd4251. PubMed PMID: 32694201; PubMed Central PMCID: PMCPMC7464562.

8. Gobeil SMC, Janowska K, McDowell S, Mansouri K, Parks R, Manne K, et al. D614G Mutation Alters SARS-CoV-2 Spike Conformation and Enhances Protease Cleavage at the S1/S2 Junction. Cell Reports. 2021;34(2):108630. doi: 10.1016/j.celrep.2020.108630.

9. Juraszek J, Rutten L, Blokland S, Bouchier P, Voorzaat R, Ritschel T, et al. Stabilizing the closed SARS-CoV-2 spike trimer. Nature Communications. 2021;12(1):244. doi: 10.1038/s41467-020-20321-x.

10. Benton DJ, Wrobel AG, Xu P, Roustan C, Martin SR, Rosenthal PB, et al. Receptor binding and priming of the spike protein of SARS-CoV-2 for membrane fusion. Nature. 2020;588(7837):327–30. Epub 20200917. doi: 10.1038/s41586-020-2772-0. PubMed PMID: 32942285; PubMed Central PMCID: PMCPMC7116727.

11. Hsieh CL, Goldsmith JA, Schaub JM, DiVenere AM, Kuo HC, Javanmardi K, et al. Structure-based design of prefusion-stabilized SARS-CoV-2 spikes. Science. 2020;369(6510):1501–5. Epub 20200723. doi: 10.1126/science.abd0826. PubMed PMID: 32703906; PubMed Central PMCID: PMCPMC7402631.

12. McCallum M, Walls AC, Bowen JE, Corti D, Veesler D. Structure-guided covalent stabilization of coronavirus spike glycoprotein trimers in the closed conformation. Nat Struct Mol Biol. 2020;27(10):942–9. Epub 20200804. doi: 10.1038/s41594-020-0483-8. PubMed PMID: 32753755; PubMed Central PMCID: PMCPMC7541350.

13. Zhou T, Tsybovsky Y, Gorman J, Rapp M, Cerutti G, Chuang GY, et al. Cryo-EM Structures of SARS-CoV-2 Spike without and with ACE2 Reveal a pH-Dependent Switch to Mediate Endosomal Positioning of Receptor-Binding Domains. Cell Host Microbe. 2020;28(6):867–79 e5. Epub 20201117. doi: 10.1016/j.chom.2020.11.004. PubMed PMID: 33271067; PubMed Central PMCID: PMCPMC7670890.

14. McCallum M, Czudnochowski N, Rosen LE, Zepeda SK, Bowen JE, Walls AC, et al. Structural basis of SARS-CoV-2 Omicron immune evasion and receptor engagement. Science. 2022;375(6583):864–8. Epub 20220125. doi: 10.1126/science.abn8652. PubMed PMID: 35076256; PubMed Central PMCID: PMCPMC9427005.

15. Zhao Z, Zhou J, Tian M, Huang M, Liu S, Xie Y, et al. Omicron SARS-CoV-2 mutations stabilize spike up-RBD conformation and lead to a non-RBM-binding monoclonal antibody escape. Nat Commun. 2022;13(1):4958. Epub 20220824. doi: 10.1038/s41467-022-32665-7. PubMed PMID: 36002453; PubMed Central PMCID: PMCPMC9399999.

16. Wrapp D, Wang N, Corbett KS, Goldsmith JA, Hsieh CL, Abiona O, et al. Cryo-EM structure of the 2019-nCoV spike in the prefusion conformation. Science. 2020;367(6483):1260–3. Epub 20200219. doi: 10.1126/science.abb2507. PubMed PMID: 32075877; PubMed Central PMCID: PMCPMC7164637.

17. Low JS, Jerak J, Tortorici MA, McCallum M, Pinto D, Cassotta A, et al. ACE2-binding exposes the SARS-CoV-2 fusion peptide to broadly neutralizing coronavirus antibodies. Science. 2022;377(6607):735–42. Epub 20220712. doi: 10.1126/science.abq2679. PubMed PMID: 35857703; PubMed Central PMCID: PMCPMC9348755.

18. Graham C, Seow J, Huettner I, Khan H, Kouphou N, Acors S, et al. Neutralization potency of monoclonal antibodies recognizing dominant and subdominant epitopes on SARS-CoV-2 Spike is impacted by the B.1.1.7 variant. Immunity. 2021;54(6):1276–89 e6. Epub 20210401. doi: 10.1016/j.immuni.2021.03.023. PubMed PMID: 33836142; PubMed Central PMCID: PMCPMC8015430.

19. Xia S, Yan L, Xu W, Agrawal AS, Algaissi A, Tseng CK, et al. A pan-coronavirus fusion inhibitor targeting the HR1 domain of human coronavirus spike. Sci Adv. 2019;5(4):eaav4580. Epub 20190410. doi: 10.1126/sciadv.aav4580. PubMed PMID: 30989115; PubMed Central PMCID: PMCPMC6457931.

20. Wild C, Oas T, McDanal C, Bolognesi D, Matthews T. A synthetic peptide inhibitor of human immunodeficiency virus replication: correlation between solution structure and viral inhibition. Proc Natl Acad Sci U S A. 1992;89(21):10537–41. doi: 10.1073/pnas.89.21.10537. PubMed PMID: 1438243; PubMed Central PMCID: PMCPMC50374.

21. Ujike M, Nishikawa H, Otaka A, Yamamoto N, Yamamoto N, Matsuoka M, et al. Heptad repeat-derived peptides block protease-mediated direct entry from the cell surface of severe acute respiratory syndrome coronavirus but not entry via the endosomal pathway. J Virol. 2008;82(1):588–92. Epub 20071017. doi: 10.1128/JVI.01697-07. PubMed PMID: 17942557; PubMed Central PMCID: PMCPMC2224400.

22. Furuta RA, Wild CT, Weng Y, Weiss CD. Capture of an early fusion-active conformation of HIV-1 gp41. Nat Struct Biol. 1998;5(4):276–9. Epub 1998/04/18. doi: 10.1038/nsb0498-276. PubMed PMID: 9546217.

23. Eckert DM, Kim PS. Mechanisms of viral membrane fusion and its inhibition. Annu Rev Biochem. 2001;70:777–810. Epub 2001/06/08. doi: 10.1146/annurev.biochem.70.1.777. PubMed PMID: 11395423.

24. Xia S, Zhu Y, Liu M, Lan Q, Xu W, Wu Y, et al. Fusion mechanism of 2019-nCoV and fusion inhibitors targeting HR1 domain in spike protein. Cell Mol Immunol. 2020;17(7):765–7. Epub 20200211. doi: 10.1038/s41423-020-0374-2. PubMed PMID: 32047258; PubMed Central PMCID: PMCPMC7075278.

25. de Vries RD, Schmitz KS, Bovier FT, Predella C, Khao J, Noack D, et al. Intranasal fusion inhibitory lipopeptide prevents direct-contact SARS-CoV-2 transmission in ferrets. Science. 2021;371(6536):1379–82. Epub 20210217. doi: 10.1126/science.abf4896. PubMed PMID: 33597220; PubMed Central PMCID: PMCPMC8011693.

26. Xia S, Liu M, Wang C, Xu W, Lan Q, Feng S, et al. Inhibition of SARS-CoV-2 (previously 2019-nCoV) infection by a highly potent pan-coronavirus fusion inhibitor targeting its spike protein that harbors a high capacity to mediate membrane fusion. Cell Res. 2020;30(4):343–55. Epub 20200330. doi: 10.1038/s41422-020-0305-x. PubMed PMID: 32231345; PubMed Central PMCID: PMCPMC7104723.

27. Hu Y, Zhu Y, Yu Y, Liu N, Ju X, Ding Q, et al. Design and characterization of novel SARS-CoV-2 fusion inhibitors with N-terminally extended HR2 peptides. Antiviral Res. 2023;212:105571. Epub 20230301. doi: 10.1016/j.antiviral.2023.105571. PubMed PMID: 36868315; PubMed Central PMCID: PMCPMC9977133.

28. Zhu Y, Yu D, Yan H, Chong H, He Y. Design of Potent Membrane Fusion Inhibitors against SARS- CoV-2, an Emerging Coronavirus with High Fusogenic Activity. J Virol. 2020;94(14). Epub 20200701. doi: 10.1128/JVI.00635-20. PubMed PMID: 32376627; PubMed Central PMCID: PMCPMC7343218.

29. Zheng M, Cong W, Peng H, Qing J, Shen H, Tang Y, et al. Stapled Peptides Targeting SARS-CoV-2 Spike Protein HR1 Inhibit the Fusion of Virus to Its Cell Receptor. J Med Chem. 2021;64(23):17486–95. Epub 20211124. doi: 10.1021/acs.jmedchem.1c01681. PubMed PMID: 34818014.

30. Yang K, Wang C, Kreutzberger AJB, Ojha R, Kuivanen S, Couoh-Cardel S, et al. Nanomolar inhibition of SARS-CoV-2 infection by an unmodified peptide targeting the prehairpin intermediate of the spike protein. Proc Natl Acad Sci U S A. 2022;119(40):e2210990119. Epub 20220919. doi: 10.1073/pnas.2210990119. PubMed PMID: 36122200; PubMed Central PMCID: PMCPMC9546559.

31. Marcink TC, Kicmal T, Armbruster E, Zhang Z, Zipursky G, Golub KL, et al. Intermediates in SARS- CoV-2 spike-mediated cell entry. Sci Adv. 2022;8(33):eabo3153. Epub 20220819. doi: 10.1126/sciadv.abo3153. PubMed PMID: 35984891; PubMed Central PMCID: PMCPMC9390989.

32. Koch J, Uckeley ZM, Doldan P, Stanifer M, Boulant S, Lozach PY. TMPRSS2 expression dictates the entry route used by SARS-CoV-2 to infect host cells. EMBO J. 2021;40(16):e107821. Epub 20210713. doi: 10.15252/embj.2021107821. PubMed PMID: 34159616; PubMed Central PMCID: PMCPMC8365257.

33. Neerukonda SN, Vassell R, Herrup R, Liu S, Wang T, Takeda K, et al. Establishment of a well-characterized SARS-CoV-2 lentiviral pseudovirus neutralization assay using 293T cells with stable expression of ACE2 and TMPRSS2. PLoS One. 2021;16(3):e0248348. Epub 20210310. doi: 10.1371/journal.pone.0248348. PubMed PMID: 33690649; PubMed Central PMCID: PMCPMC7946320.

34. Buchrieser J, Dufloo J, Hubert M, Monel B, Planas D, Rajah MM, et al. Syncytia formation by SARS-CoV-2-infected cells. EMBO J. 2020;39(23):e106267. Epub 20201104. doi: 10.15252/embj.2020106267. PubMed PMID: 33051876; PubMed Central PMCID: PMCPMC7646020.

35. Zeng C, Evans JP, King T, Zheng YM, Oltz EM, Whelan SPJ, et al. SARS-CoV-2 spreads through cell-to-cell transmission. Proc Natl Acad Sci U S A. 2022;119(1):e2111400119. doi: 10.1073/pnas.2111400119. PubMed PMID: 34937699; PubMed Central PMCID: PMCPMC8740724.

36. Papa G, Mallery DL, Albecka A, Welch LG, Cattin-Ortola J, Luptak J, et al. Furin cleavage of SARS- CoV-2 Spike promotes but is not essential for infection and cell-cell fusion. PLoS Pathog. 2021;17(1):e1009246. Epub 20210125. doi: 10.1371/journal.ppat.1009246. PubMed PMID: 33493182; PubMed Central PMCID: PMCPMC7861537.

37. Reuter N, Chen X, Kropff B, Peter AS, Britt WJ, Mach M, et al. SARS-CoV-2 Spike Protein Is Capable of Inducing Cell&ndash;Cell Fusions Independent from Its Receptor ACE2 and This Activity Can Be Impaired by Furin Inhibitors or a Subset of Monoclonal Antibodies. Viruses. 2023;15(7):1500. PubMed PMID: doi:10.3390/v15071500.

38. Yeung ML, Teng JLL, Jia L, Zhang C, Huang C, Cai JP, et al. Soluble ACE2-mediated cell entry of SARS-CoV-2 via interaction with proteins related to the renin-angiotensin system. Cell. 2021;184(8):2212–28 e12. Epub 20210302. doi: 10.1016/j.cell.2021.02.053. PubMed PMID: 33713620; PubMed Central PMCID: PMCPMC7923941.

39. He Y, Vassell R, Zaitseva M, Nguyen N, Yang Z, Weng Y, et al. Peptides trap the human immunodeficiency virus type 1 envelope glycoprotein fusion intermediate at two sites. J Virol. 2003;77(3):1666–71. Epub 2003/01/15. doi: 10.1128/jvi.77.3.1666-1671.2003. PubMed PMID: 12525600; PubMed Central PMCID: PMCPMC140873.

40. De Feo CJ, Wang W, Hsieh ML, Zhuang M, Vassell R, Weiss CD. Resistance to N-peptide fusion inhibitors correlates with thermodynamic stability of the gp41 six-helix bundle but not HIV entry kinetics. Retrovirology. 2014;11:86. Epub 20141002. doi: 10.1186/s12977-014-0086-8. PubMed PMID: 25274545; PubMed Central PMCID: PMCPMC4190581.

41. Golding H, Zaitseva M, de Rosny E, King LR, Manischewitz J, Sidorov I, et al. Dissection of human immunodeficiency virus type 1 entry with neutralizing antibodies to gp41 fusion intermediates. J Virol. 2002;76(13):6780–90. doi: 10.1128/jvi.76.13.6780-6790.2002. PubMed PMID: 12050391; PubMed Central PMCID: PMCPMC136262.

42. Du L, Kao RY, Zhou Y, He Y, Zhao G, Wong C, et al. Cleavage of spike protein of SARS coronavirus by protease factor Xa is associated with viral infectivity. Biochem Biophys Res Commun. 2007;359(1):174–9. Epub 20070522. doi: 10.1016/j.bbrc.2007.05.092. PubMed PMID: 17533109; PubMed Central PMCID: PMCPMC2323977.

43. Lavie M, Dubuisson J, Belouzard S. SARS-CoV-2 Spike Furin Cleavage Site and S2’ Basic Residues Modulate the Entry Process in a Host Cell-Dependent Manner. J Virol. 2022;96(13):e0047422. Epub 20220609. doi: 10.1128/jvi.00474-22. PubMed PMID: 35678602; PubMed Central PMCID: PMCPMC9278140.

44. Yu S, Zheng X, Zhou B, Li J, Chen M, Deng R, et al. SARS-CoV-2 spike engagement of ACE2 primes S2’ site cleavage and fusion initiation. Proc Natl Acad Sci U S A. 2022;119(1):e2111199119. doi: 10.1073/pnas.2111199119. PubMed PMID: 34930824; PubMed Central PMCID: PMCPMC8740742.

45. Shi R, Shan C, Duan X, Chen Z, Liu P, Song J, et al. A human neutralizing antibody targets the receptor-binding site of SARS-CoV-2. Nature. 2020;584(7819):120–4. Epub 20200526. doi: 10.1038/s41586-020-2381-y. PubMed PMID: 32454512.

46. Barnes CO, Jette CA, Abernathy ME, Dam KA, Esswein SR, Gristick HB, et al. SARS-CoV-2 neutralizing antibody structures inform therapeutic strategies. Nature. 2020;588(7839):682–7. Epub 20201012. doi: 10.1038/s41586-020-2852-1. PubMed PMID: 33045718; PubMed Central PMCID: PMCPMC8092461.

47. Westendorf K, Zentelis S, Wang L, Foster D, Vaillancourt P, Wiggin M, et al. LY-CoV1404 (bebtelovimab) potently neutralizes SARS-CoV-2 variants. Cell Rep. 2022;39(7):110812. Epub 20220425. doi: 10.1016/j.celrep.2022.110812. PubMed PMID: 35568025; PubMed Central PMCID: PMCPMC9035363.

48. Du X, Tang H, Gao L, Wu Z, Meng F, Yan R, et al. Omicron adopts a different strategy from Delta and other variants to adapt to host. Signal Transduct Target Ther. 2022;7(1):45. Epub 20220210. doi: 10.1038/s41392-022-00903-5. PubMed PMID: 35145066; PubMed Central PMCID: PMCPMC8830988.

49. Meng B, Abdullahi A, Ferreira I, Goonawardane N, Saito A, Kimura I, et al. Altered TMPRSS2 usage by SARS-CoV-2 Omicron impacts infectivity and fusogenicity. Nature. 2022;603(7902):706–14. Epub 20220201. doi: 10.1038/s41586-022-04474-x. PubMed PMID: 35104837; PubMed Central PMCID: PMCPMC8942856.

50. Willett BJ, Grove J, MacLean OA, Wilkie C, De Lorenzo G, Furnon W, et al. SARS-CoV-2 Omicron is an immune escape variant with an altered cell entry pathway. Nature Microbiology. 2022;7(8):1161–79. doi: 10.1038/s41564-022-01143-7.

51. Gobeil SM, Henderson R, Stalls V, Janowska K, Huang X, May A, et al. Structural diversity of the SARS-CoV-2 Omicron spike. Mol Cell. 2022;82(11):2050–68 e6. Epub 20220325. doi: 10.1016/j.molcel.2022.03.028. PubMed PMID: 35447081; PubMed Central PMCID: PMCPMC8947964.

52. Nishima W, Kulik M. Full-Length Computational Model of the SARS-CoV-2 Spike Protein and Its Implications for a Viral Membrane Fusion Mechanism. Viruses. 2021;13(6). Epub 20210611. doi: 10.3390/v13061126. PubMed PMID: 34208191; PubMed Central PMCID: PMCPMC8230804.

53. Costello SM, Shoemaker SR, Hobbs HT, Nguyen AW, Hsieh CL, Maynard JA, et al. The SARS-CoV-2 spike reversibly samples an open-trimer conformation exposing novel epitopes. Nat Struct Mol Biol. 2022;29(3):229–38. Epub 20220302. doi: 10.1038/s41594-022-00735-5. PubMed PMID: 35236990; PubMed Central PMCID: PMCPMC9007726.

54. Raghuvamsi PV, Tulsian NK, Samsudin F, Qian X, Purushotorman K, Yue G, et al. SARS-CoV-2 S protein:ACE2 interaction reveals novel allosteric targets. Elife. 2021;10. Epub 20210208. doi: 10.7554/eLife.63646. PubMed PMID: 33554856; PubMed Central PMCID: PMCPMC7932696.

55. Kobayashi J, Kanou K, Okura H, Akter TM, Fukushi S, Matsuyama S. Biochemical analysis of packing and assembling heptad repeat motifs in the coronavirus spike protein trimer. mBio. 2024:e0120324. Epub 20241023. doi: 10.1128/mbio.01203-24. PubMed PMID: 39440974.

56. Casalino L, Gaieb Z, Goldsmith JA, Hjorth CK, Dommer AC, Harbison AM, et al. Beyond Shielding: The Roles of Glycans in the SARS-CoV-2 Spike Protein. ACS Cent Sci. 2020;6(10):1722–34. Epub 20200923. doi: 10.1021/acscentsci.0c01056. PubMed PMID: 33140034; PubMed Central PMCID: PMCPMC7523240.

57. Zoufaly A, Poglitsch M, Aberle JH, Hoepler W, Seitz T, Traugott M, et al. Human recombinant soluble ACE2 in severe COVID-19. The Lancet Respiratory Medicine. 2020;8(11):1154–8. doi: 10.1016/S2213-2600(20)30418-5.

58. Brandenberg OF, Magnus C, Rusert P, Regoes RR, Trkola A. Different Infectivity of HIV-1 Strains Is Linked to Number of Envelope Trimers Required for Entry. PLOS Pathogens. 2015;11(1):e1004595. doi: 10.1371/journal.ppat.1004595.

59. Yang X, Kurteva S, Ren X, Lee S, Sodroski J. Stoichiometry of Envelope Glycoprotein Trimers in the Entry of Human Immunodeficiency Virus Type 1. Journal of Virology. 2005;79(19):12132–47. doi: doi:10.1128/jvi.79.19.12132-12147.2005.

60. Crawford KHD, Eguia R, Dingens AS, Loes AN, Malone KD, Wolf CR, et al. Protocol and Reagents for Pseudotyping Lentiviral Particles with SARS-CoV-2 Spike Protein for Neutralization Assays. Viruses. 2020;12(5):513. PubMed PMID: doi:10.3390/v12050513.

61. Lee SH, Mayr C. Gain of Additional BIRC3 Protein Functions through 3’-UTR-Mediated Protein Complex Formation. Mol Cell. 2019;74(4):701–12.e9. Epub 20190401. doi: 10.1016/j.molcel.2019.03.006. PubMed PMID: 30948266; PubMed Central PMCID: PMCPMC6581197.

